## Supplementary Material for "β-carotene accelerates the resolution of atherosclerosis in mice"

**Supplementary Figures**

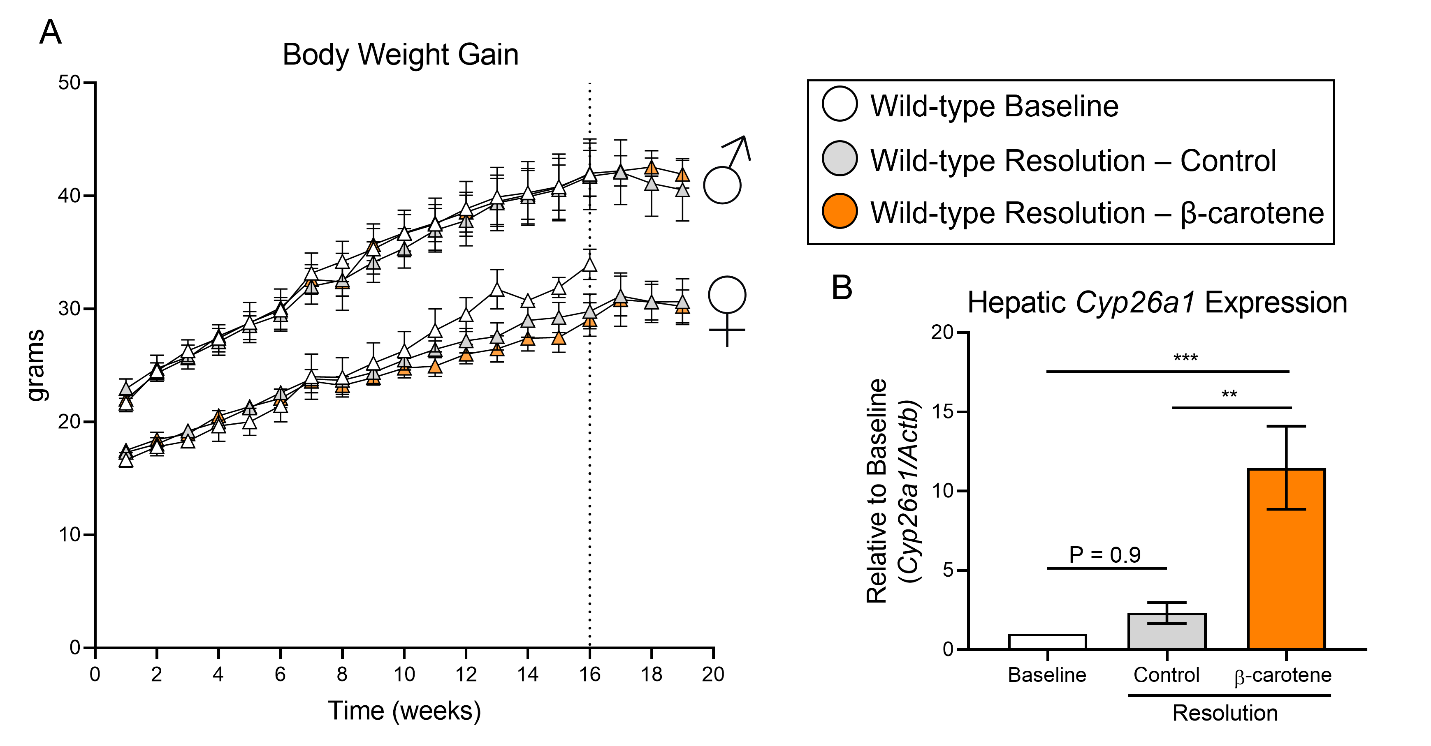

**Supplementary Figure 1.** Four-week-old male and female wild-type mice were fed a purified Western diet deficient in vitamin A (WD-VAD) and injected with antisense oligonucleotide targeting the low-density lipoprotein receptor (ASO-LDLR) once a week for 16 weeks to induce atherosclerosis. After 16 weeks (dotted line), a group of mice was harvested (Baseline) and the rest of the mice were injected once with sense oligonucleotide (SO-LDLR) to inactivate ASO-LDLR and promote atherosclerosis resolution. Mice undergoing resolution were either kept on the same diet (Resolution - Control) or switched to a Western diet supplemented with 50 mg/kg of β-carotene (Resolution - β-carotene) for three more weeks**. (A**) body weight progression, and **(B)** hepatic expression mRNA expression for *Cyp26a1* refered to *Actb* as a houskeeping control. N = 5 to 10 mice/group, Values are represented as means ± SEM. Statistical differences were evaluated using one-way ANOVA with Tukey’s multiple comparisons test. Differences between groups were considered significant with a p-value < 0.05. ** p < 0.01; *** p < 0.005.

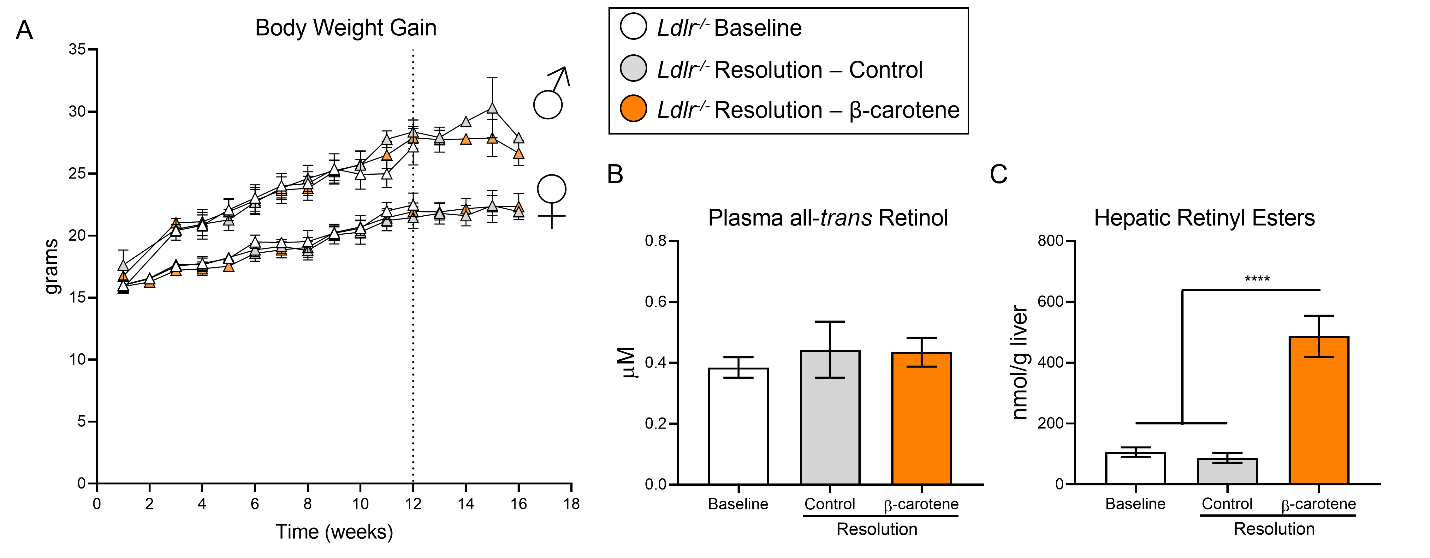

**Supplementary Figure 2.** Four-week-old male and female *Ldlr^-/-^* mice were fed a purified Western diet deficient in vitamin A (WD-VAD) for 12 weeks to induce atherosclerosis. After 12 weeks (dotted line), a group of mice was harvested (Baseline) and the rest of the mice were switched to a Standard diet (Resolution-Control) or the same diet supplemented with 50 mg/kg of β-carotene (Resolution-β-carotene) for four more weeks. **(A**) body weight progression. **(B)** circulating vitamin A (all-*trans* retinol), and **(C)** hepatic retinyl ester stores. N = 5 to 10 mice/group, Values are represented as means ± SEM. Statistical differences were evaluated using one-way ANOVA with Tukey’s multiple comparisons test. Differences between groups were considered significant with a p-value < 0.05. **** p < 0.001.

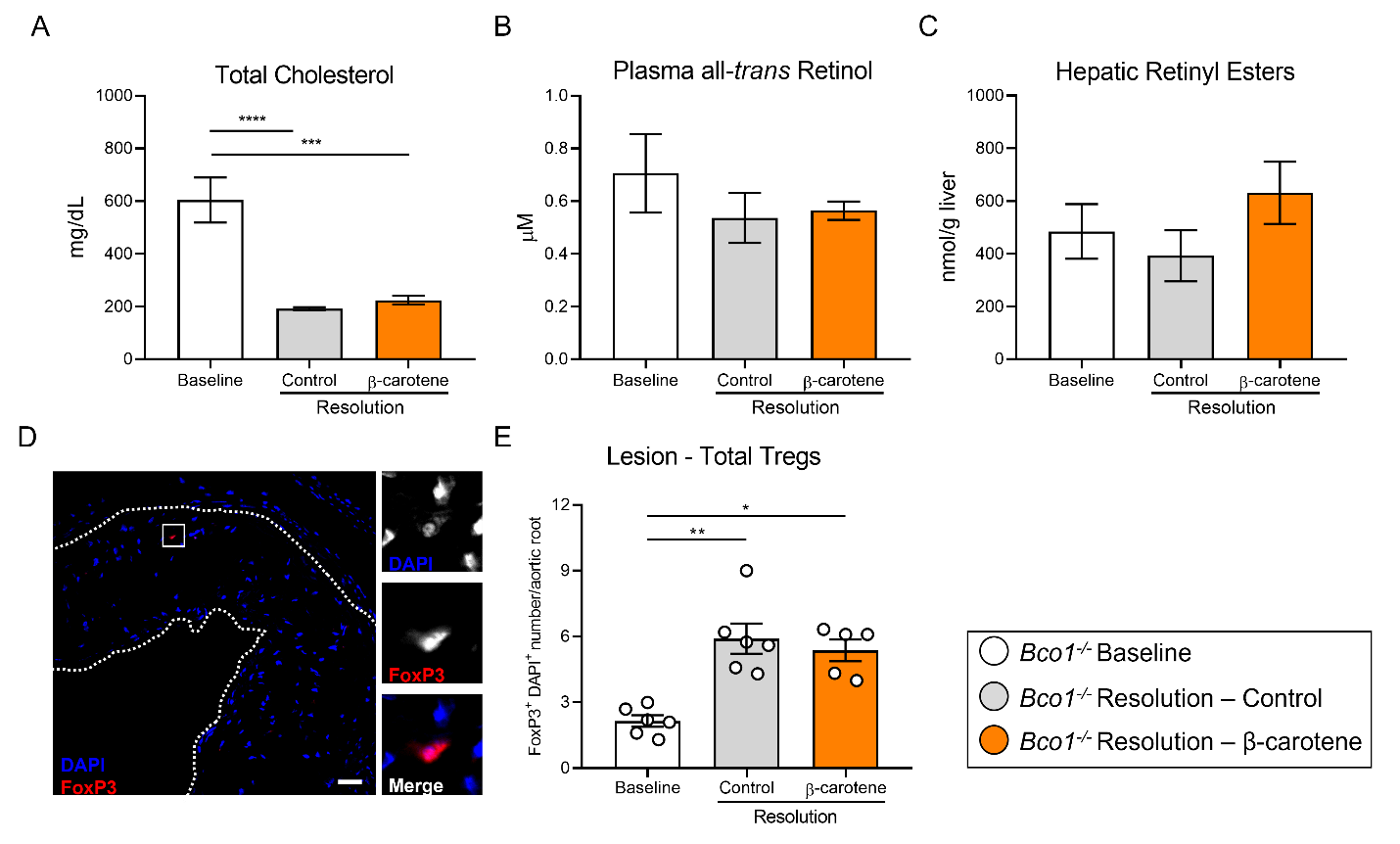

**Supplementary Figure 3.** Four-week-old male and female *Bco1^-/-^* mice were fed a purified Western diet deficient in vitamin A (WD-VAD) and injected with antisense oligonucleotide targeting the low-density lipoprotein receptor (ASO-LDLR) once a week for 16 weeks to induce the development of atherosclerosis. After 16 weeks, a group of mice was harvested (Baseline) and the rest of the mice were injected once with sense oligonucleotide (SO-LDLR) to block ASO-LDLR (Resolution). Mice undergoing resolution were either kept on the same diet (Resolution-Control) or switched to a Western diet supplemented with 50 mg/kg of β-carotene (Resolution- β-carotene) for three more weeks. **(A)** Total plasma cholesterol, and **(B)** vitamin A (all-*trans* retinol). **(C)** Hepatic vitamin A (retinyl ester) stores. **(D)** Representative confocal image and **(E)** quantification of total Tregs in the lesion. Size bar 50 µm. N = 5 to 10 mice/group, Values are represented as means ± SEM. Statistical differences were evaluated using one-way ANOVA with Tukey’s multiple comparisons test. Differences between groups were considered significant with a p-value < 0.05. *** p < 0.005, **** p < 0.001.

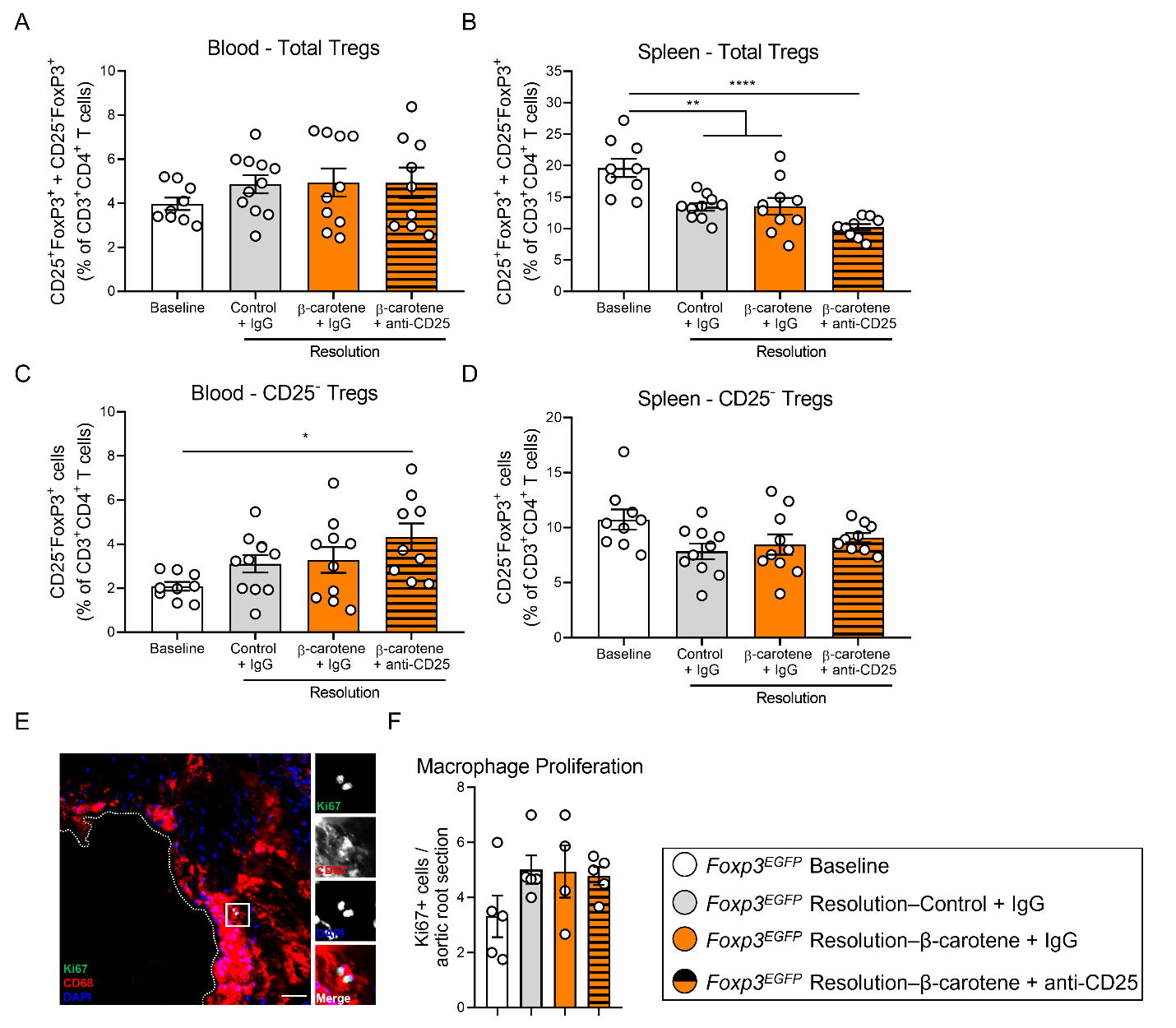

**Supplementary Figure 4.** Four-week-old male and female mice expressing enhanced green fluorescence protein (EGFP) under the control of the forkhead box P3 (*Foxp3*) promoter (*Foxp3^EGFP^* mice) were fed a purified Western diet deficient in vitamin A (WD-VAD) and injected with antisense oligonucleotide targeting the low-density lipoprotein receptor (ASO-LDLR) once a week for 16 weeks to induce the development of atherosclerosis. After 16 weeks, a group of mice was harvested (Baseline) and the rest of the mice were injected once with sense oligonucleotide (SO-LDLR) to block ASO-LDLR (Resolution). Mice undergoing resolution were either kept on the same diet (Resolution-Control) or switched to a Western diet supplemented with 50 mg/kg of β-carotene (Resolution- β-carotene) for three more weeks. An additional group of mice fed with β-carotene was injected twice before sacrifice with anti-CD25 monoclonal antibody to deplete Treg (Resolution-β-carotene+anti-CD25). The rest of the resolution groups were injected with IgG isotype control antibody. **(A)** Quantification of the circulating and **(B)** splenic CD25^+^FoxP3^+^ and CD25^-^FoxP3^+^ (Total Tregs) measured by flow cytometry. **(C)** Circulating and **(D)** splenic CD25^-^FoxP3^+^ Tregs. **(E)** Macrophages proliferating in the lesion were identified by the colocalization of Ki67 (green) and DAPI (blue) in CD68^+^ (red) cells. **(F)** Number of Ki67+ macrophages in the lesion. Size bars = 50 μm. N = 4 to 10 mice/group. Values are represented as means ± SEM. Statistical differences were evaluated using one-way ANOVA with Tukey’s multiple comparisons test. Differences between groups were considered significant with a p-value < 0.05. * p < 0.05.

**Supplementary Table 1.** Composition of the experimental diets utilized in the study.

| **Ingredient** | **WD-**  **VAD**  **(g/kg diet)** | **WD-**  **β-carotene**  **(g/kg diet)** | **Standard-**  **VAD**  **(g/kg diet)** | **Standard-**  **β-carotene**  **(g/kg diet)** |
| --- | --- | --- | --- | --- |
| **Casein** | 200 | 200 | 200 | 200 |
| **L-Cysteine** | 3 | 3 | 3 | 3 |
| **Corn starch** | 72.8 | 72.8 | 319 | 319 |
| **Maltodextrin** | 100 | 100 | 100 | 100 |
| **Sucrose** | 212 | 212 | 212 | 212 |
| **Cellulose** | 50 | 50 | 50 | 50 |
| **Soybean oil** | 25 | 25 | 70 | 70 |
| **Lard** | 160 | 160 | 0 | 0 |
| **t-Butylhydroquinone** | 0 | 0 | 0 | 0 |
| **Choline bitartrate** | 2 | 2 | 2 | 2 |
| **Dicalcium phosphate** | 13 | 13 | 13 | 13 |
| **Calcium carbonate** | 5.5 | 5.5 | 5.5 | 5.5 |
| **Potassium citrate monohydrate** | 16.5 | 16.5 | 16.5 | 16.5 |
| **Cholesterol** | 3.08 | 3.08 | 0 | 0 |
| **Mineral mix** | 10 | 10 | 35 | 35 |
| **Vitamin mix, no added vitamin A** | 10 | 10 | 10 | 10 |
| **Placebo beadlets** | 0.5 | 0 | 0.5 | 0 |
| **β-carotene beadlets, 10% β-carotene** | 0 | 0.5 | 0 | 0.5 |

### ^1^ Footnotes. IU: International unit. WD, Western diet; VAD; Vitamin A deficient; IU, international units.
